# Three-dimensional cultured Liver-on-a-Chip with mature hepatocyte-like cells derived from human pluripotent stem cells

**DOI:** 10.1101/232215

**Authors:** Ken-ichiro Kamei, Momoko Yoshioka, Shiho Terada, Yumie Tokunaga, Yong Chen

## Abstract

Liver-on-a-Chip technology holds considerable potential for applications in drug screening and chemical-safety testing. To establish such platforms, functional hepatocytes are required; however, primary hepatocytes are commonly used, despite problems involving donor limitations, lot-to-lot variation, and unsatisfactory two-dimensional culture methods. Although human pluripotent stem cells (hPSCs) may represent a strong alternative contender to address the aforementioned issues, remaining technological challenges include the robust, highly efficient production of high-purity hepatic clusters. In addition, current Liver-on-a-Chip platforms are relatively complicated and not applicable for high-throughput experiments. Here, we develop a very simple Liver-on-a-Chip platform with mature and functional hepatocyte-like cells derived from hPSCs. To establish a method for hepatic differentiation of hPSCs, cells were first treated by inhibiting the phosphoinositide 3-kinase- and Rho-associated protein kinase-signaling pathways to stop self-renewal and improve survival, respectively, which enabled the formation of a well-defined endoderm and facilitated hepatocyte commitment. Next, a simple microfluidic device was used to create a three-dimensional (3D) culture environment that enhanced the maturation and function of hepatocyte-like cells by increasing the expression of both hepatic maturation markers and cytochrome P450. Finally, we confirmed improvements in hepatic functions, such as drug uptake/excretion capabilities, in >90% of 3D-matured hepatocyte-like cells by indocyanin green assay. These results indicated that the incorporation of hPSC-derived hepatocytes on our Liver-on-a-Chip platform may serve to enhance the processes involved in drug screening and chemical-safety testing.

## Introduction

Organs-on-a-Chip (OoC) platforms hold considerable potential for pre-clinical trials of drug development^1–3^ as well as toxicological tests.^4^ To date, such testing has relied upon animal models; however, large challenges remain with regard to predicting the safety and efficacy of drug candidates for human clinical use. Therefore, new *in vitro* models need to be developed as an alternative to animal models to predict treatment effects on humans with high predictability. OoCs represent the most suitable platforms to fulfill these requirements, as these platforms were developed to recapitulate human physiological conditions *in vitro*. Currently, numerous OoC platforms have been reported as representative of organs, such as the brain,^5^ lung,^6,7^ heart,^8,9^ gut,^10,11^ intestine,^12^ liver^13–16^ or multiple tissues.^17–19^ Considering that the liver constitutes the largest organ in the human body and has many critical roles for body maintenance as well as drug metabolism and detoxification, we focused on optimizing the Liver-on-a-Chip platform, as the reported platform still retains two critical issues involving cell sources and cell culture methods.

In terms of cell source, hepatocytes, which constitute the majority of cells in the liver, are mainly used for current drug screening and toxicological testing as well as Liver-on-a-Chip platforms. Primary hepatocytes can be obtained from healthy donors; however, these cells cannot proliferate under standard cell-culture conditions and often experience loss of function. As an alternative, established cell lines are used for cell culture and *in vitro* drug testing, although these cells were often originally harvested from tumors and do not represent healthy liver functions. Therefore, obtaining sufficient amounts of proper and functional hepatocytes remains challenging.

To meet this requirement, human pluripotent stem cells (hPSCs), such as embryonic (hESCs^20^) and induced pluripotent stem cells (hiPSCs^21^) hold considerable potential as sources to generate hepatocytes. hPSCs exhibit two distinctive capabilities: unlimited self-renewal without karyotypical abnormality and the ability to differentiate into almost any cell type. These capabilities allow for the generation of hepatocyte quantities sufficient for subsequent applications such as Liver-on-a-Chip platforms.

However, there remain issues with current methods related to the induction of hepatic differentiation from hPSCs to result in functional and mature hepatocytes. These methods involve either 1) formation of embryoid bodies or cell aggregates,^22^ or 2) introduction of exogenous genes (e.g., *HHEX*^23^ and *SOX17*^24^). Such methods have high potential to cause quality control difficulties involving contamination with other differentiated cells; moreover, the hepatic functionality of the resultant cells still has considerable room for improvement. Therefore, it is necessary to establish a novel method enabling the efficient and robust differentiation of hPSCs into functional hepatocytes.

In addition to cell sources, cell culture methods also require new innovation. Although two-dimensional (2D) culture of hepatocytes has been carried out for many years, this method is also problematic; for example, primary hepatocytes often lose their function under 2D culture. Therefore, a number of reports have suggested the use of three-dimensional (3D) culture methods to render hepatocytes more functional.^25^ Moreover, recent studies have shown that organoids^26–28^ hold promise to obtain micro live tissues with greater functionality than those of 2D culture. However, the described cell aggregates and organoids exhibited varying size and structure as they are reliant upon only the capacity of the stem cells for tissue formation, resulting in low reproducibility. Therefore, these methods have yet to reach an acceptable stage for use in drug screening. Similarly, the majority of conventional Liver-on-a-Chip platforms were based on 2D cell culture. Moreover, as most previously developed Liver-on-a-Chip platforms are both sophisticated and complicated and require special instruments to control the procedures, they are not user-friendly, limiting their use by general biology laboratories and their applicability for drug screening owing to the lack of high-throughput screening capability.

Here, we developed a novel Liver-on-a-Chip platform by 1) establishing a protocol enabling the efficient differentiation of hepatocyte-like cells from hPSCs, resulting in high degrees of purity and reproducibility, and 2) applying a simple microfluidic 3D cell-culture platform^29,30^ (**Fig. 1**). To facilitate efficient hepatic differentiation and cell survival, our method utilizes a cocktail of chemicals and growth factors. Conversely, our method does not require the use of exogenous genes, thereby eliminating concerns of genetic integration into host cells and variations in gene-delivery efficiency. After hepatic differentiation from hPSCs to hepatic progenitor cells, the cells are introduced into our simple microfluidic 3D cell culture device to prepare our Liver-on-a-Chip platform with hPSC-derived hepatocyte-like cells of high-maturation status. Notably, this platform does not require any special instruments, with standard pipettes able to be used to carry out the experiments.

**Fig. 1.**
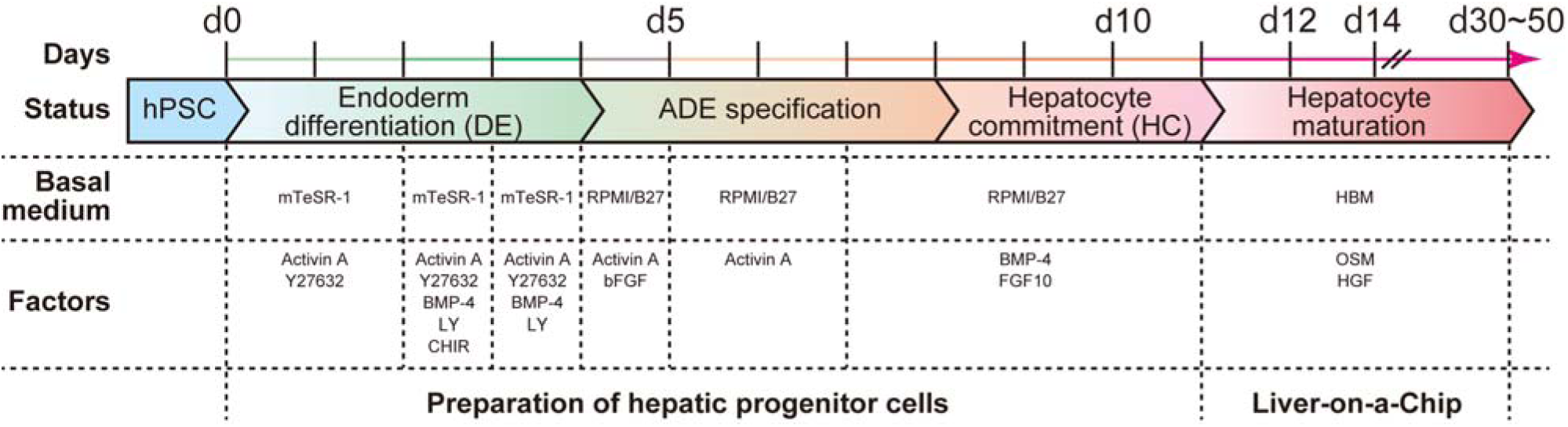
The hepatic differentiation process from human pluripotent stem cells. A variety of solutions containing basal medium, growth factors, and chemicals were administered to hPSCs to efficiently induce differentiation to hepatocyte-like cells. Initially, hPSCs were treated with mTeSR-1 medium supplemented with combinations of Activin A, BMP-4, CHIR99021 (CHIR), LY294002 (LY), and Y27632 (Y) to induce definitive endoderm (DE) differentiation until Day 4 (d4). Then, cells were treated with Activin A in RPMI medium supplemented with B27 for anterior definitive endoderm (ADE) specification until Day 8. As the next step, cells were treated with BMP-4 and FGF-10 for inducing hepatocyte commitment in RPMI medium supplemented with B27 until Day 11. Finally, cells were treated with oncostatin M (OSM) and hepatocyte growth factor (HGF) in hepatocyte basal medium for hepatocyte maturation.

## Results and discussion

### Cell survival and robustness of definitive-endoderm (DE) differentiation is enhanced by phosphoinositide 3-kinase (PI3K) and Rho-associated protein kinase (ROCK) inhibition

Induction of DE differentiation is the most critical step required to promote further differentiation (**Fig. 2**). In this process, transforming growth factor (TGF)-α,^31,32^ bone morphogenetic protein (BMP)-4,^33^ α-catenin,^34,35^ and basic fibroblast growth factor (bFGF)^32,36^-signaling pathways play critical roles. Moreover, the PI3K-signaling pathway is critical for the maintenance of hPSC self-renewal,^37,38^ with inhibition of this pathway resulting in hPSC differentiation. Previously, Hannan et al. established a method to effectively achieve hPSC differentiation to DE using a cocktail of activin A, BMP-4, bFGF, LY294002, and CHIR99021, but without the use of exogenous genes.^39^ However, this method results in massive cell death and inefficient and irreproducible outcomes. Furthermore, the use of hPSC aggregates is unlikely to allow improved robustness, owing to the challenge associated with size control, which is a critical factor for lineage determination during hPSC differentiation. Although the use of dissociated hPSCs may improve robustness, the related processes continue to result in large amounts of cell death. Therefore, we used a ROCK inhibitor (Y27632) that has previously been used to facilitate the survival of dissociated, self-renewing hPSCs.^40^ We hypothesized that administration of a ROCK inhibitor would also facilitate cell survival during the early stage of DE differentiation. For the medium, we modified mTeSR-1-defined medium^41^ containing BMP- and bFGF (**Fig. 1** and **Supplementary Table S1**), which are also commonly used as key factors to introduce DE differentiation. We observed that hPSCs showed unstable behavior following introduction of the new culture conditions; therefore, for establishing robust differentiation protocols, the initial use of self-renewing culture medium was beneficial, followed by gradually changing to the differentiation medium. Toward this end, activin A, BMP-4, LY294002, and CHIR99021 were subsequently added into the mTeSR-1 medium for DE differentiation.

**Fig. 2.**
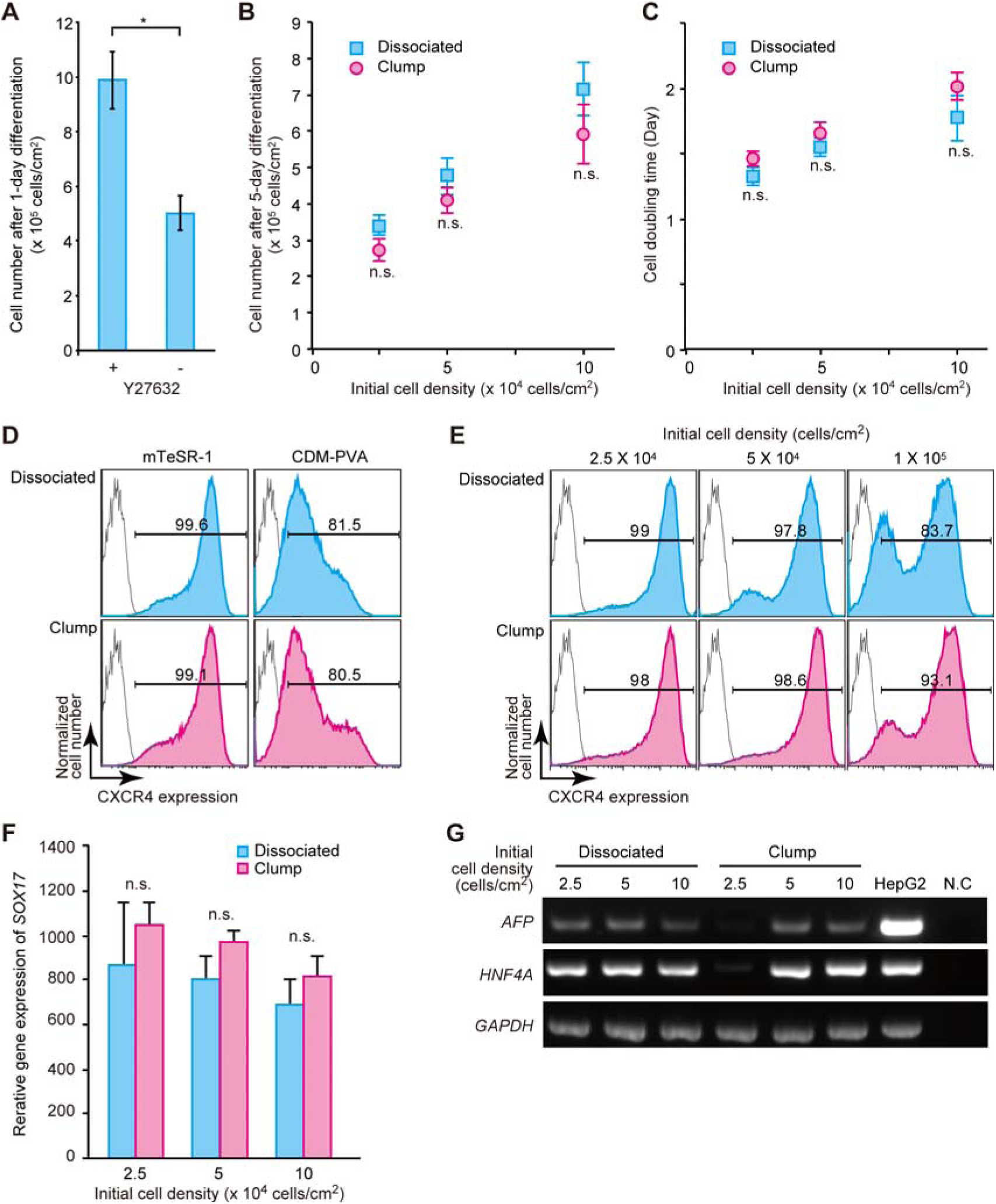
Combination of ROCK (10 µM Y27632) and PI3K (10 µM LY294002) inhibitors facilitates survival and inhibits self-renewal of hPSCs during definitive-endoderm (DE) differentiation. (**A**) 1-day treatment with mTeSR-1 basal medium in combination with Y27632 increases cell survival of H9 human embryonic stem cells (hESCs). Data represent the means ± standard deviation (*n* = 3). (**B, C**) Both dissociated and clumped H9 hESCs treated with mTeSR-1/Y27632 were able to proliferate during DE differentiation (**B**) with similar doubling times (**C**). Data represent the means ± standard deviation (*n* = 3). “n.s.” represents “not significant”. (**D**) Selection of mTeSR-1 or CDM-PVA medium to induce differentiation of dissociated or clumped H9 hESCs into DE by monitoring the expression levels of the CXCR4 DE marker. Cells treated with mTeSR-1 medium supplemented with activin A, BMP-4, LY294002, and CHIR99021, along with Y27632 showed a more homogeneous cell population with higher CXCR4 expression. (**E**) Initial cell density influences the cellular heterogeneity during DE differentiation, as determined by CXCR4 expression. (**F**) Initial cell density is inversely correlated with expression levels of the *SOX17* DE marker. At low initial cell density, *SOX17* expression levels varied among experiments. Data represent the means ± standard deviation (*n* = 3). “n.s.” represents “not significant”. (**G**) Representative gel electrophoresis of RT-PCR samples to ascertain expression of α-fetoprotein (*AFP*) and hepatocyte nuclear factor 4α (*HNF4A*) differentiation marker expression on day 12. *GAPDH*, a housekeeping gene, was used as an internal control. Data were obtained from three independent experiments. N.C. represents a negative control (water).

To confirm increases in cell survival during DE differentiation, we counted H9 hESCs^20^ following 1 day of treatment (**Fig. 2A**). Cells treated with both 10 µM Y27632 and 10 µM LY294002 showed significant increases in cell number, as compared with levels observed in cells treated with only LY294002. This result indicated that inhibition of the ROCK-signaling pathway improved cell survival to a degree greater than that observed following inhibition of the PI3K-signaling pathway.

We then investigated differences in outcomes based on initiating differentiation from either dissociated or clumped hPSCs according to cell number and doubling time (**Fig. 2B and C**). We observed that following a 5-day incubation with 10 µM Y27632 and 10 µM LY294002 and DE differentiation, both dissociated and clumped hESCs increased cell numbers with similar doubling times. These results suggested that Y27632 improves cell survival even during DE differentiation, but not cell proliferation.

To confirm cell status following DE differentiation, the expression of C-X-C-motif chemokine receptor-4 (CXCR4), a cell surface DE-differentiation marker, was quantitatively analyzed by flow cytometry (**Fig. 2D**) and compared with levels measured in previously-reported chemically defined medium (CDM-PVA)^39^ (**Supplementary Table S2**). Both H9 and 253G1 cells treated with mTeSR-based medium showed higher degrees of purity and enhanced CXCR4 expression (99.6%) compared to those of cells cultured in CDM-PVA medium (81.5%). Then, we investigated the effects of initial cell-seeding density on the induction of DE differentiation (**Fig. 2E**). During 5-day DE differentiation, cell numbers in all cell densities increased approximately 10 fold (**Fig. 2B**); however, the increase in initial cell number revealed bimodal cell populations with regard to CXCR4 expression (**Fig. 2E**). These results indicated that the use of a smaller initial cell-seeding density resulted in a more homogeneous cell population with higher CXCR4 expression.

We also measured expression of the sex-determining region Y box-17 (*SOX17*) transcription factor and endoderm-lineage marker by quantitative reverse transcription-polymerase chain reaction (RT-PCR) (**Fig. 2F**). The use of cell clumps resulted in higher *SOX17* expression as compared with using dissociated cells, with *SOX17* expression subsequently decreasing along with increasing cell numbers, although lower initial cell-seeding density (2.5 × 10^4^ cells cm^−2^) resulted in large variations in *SOX17* expression. We also confirmed hepatocyte differentiation by monitoring the expression of α-fetoprotein (*AFP*) and hepatocyte nuclear factor 4α (*HNF4A* differentiation marker mRNA on day 12 (**Fig. 2G**). The use of cell aggregates with lower initial cell-seeding density (2.5 × 10^4^ cells cm^−2^) did not result in *AFP* and *HNF4A* expression. Based on these results, we used 5 × 10^4^ cells cm^−2^ for further differentiation protocols consequent to our observation of higher purity and improved robustness at higher initial cell densities. Under the optimized conditions, we tested DE differentiation from 253G1 hiPSCs^42^ and differentiation to highly pure CXCR4-positive DE cells, and found that this condition gave similar results as H9 hESCs, even with 253G1 hiPSCs (**Supplementary Fig. S1**).

### Optimization to induce anterior definitive-endoderm (ADE) specification and hepatocyte commitment (HC)

We established a robust and efficient procedure for ADE specification from DE cells, as well as HCs, to generate hepatic progenitor cells (**Fig. 3** and **Supplementary Table S1**). For ADE specification as shown in **Fig. 1** and **Experimental**, it was necessary to maintain activin A levels but remove bFGF to reduce active cell-signaling pathways and maintain pluripotency.^43^ Alternatively, HC requires BMP4 and FGF10 signaling.^44^ ADE specification and HC require several days to achieve treatment-induced changes in phenotype. Therefore, we treated cells with activin A and a combination of BMP4 and FGF10 in RPMI medium supplemented with B27 supplement for ADE and HC, respectively, and varied these treatment periods to identify the optimal timing (~1–3 days for ADE and ~1–4 days for HC) for hepatic progenitor cell differentiation (**Fig. 3A**) by observing the expression levels of genes associated with each differentiation phenotype following treatment as determined by quantitative RT-PCR (**Fig. 3B** and **Supplementary Table S3**).

**Fig. 3.**
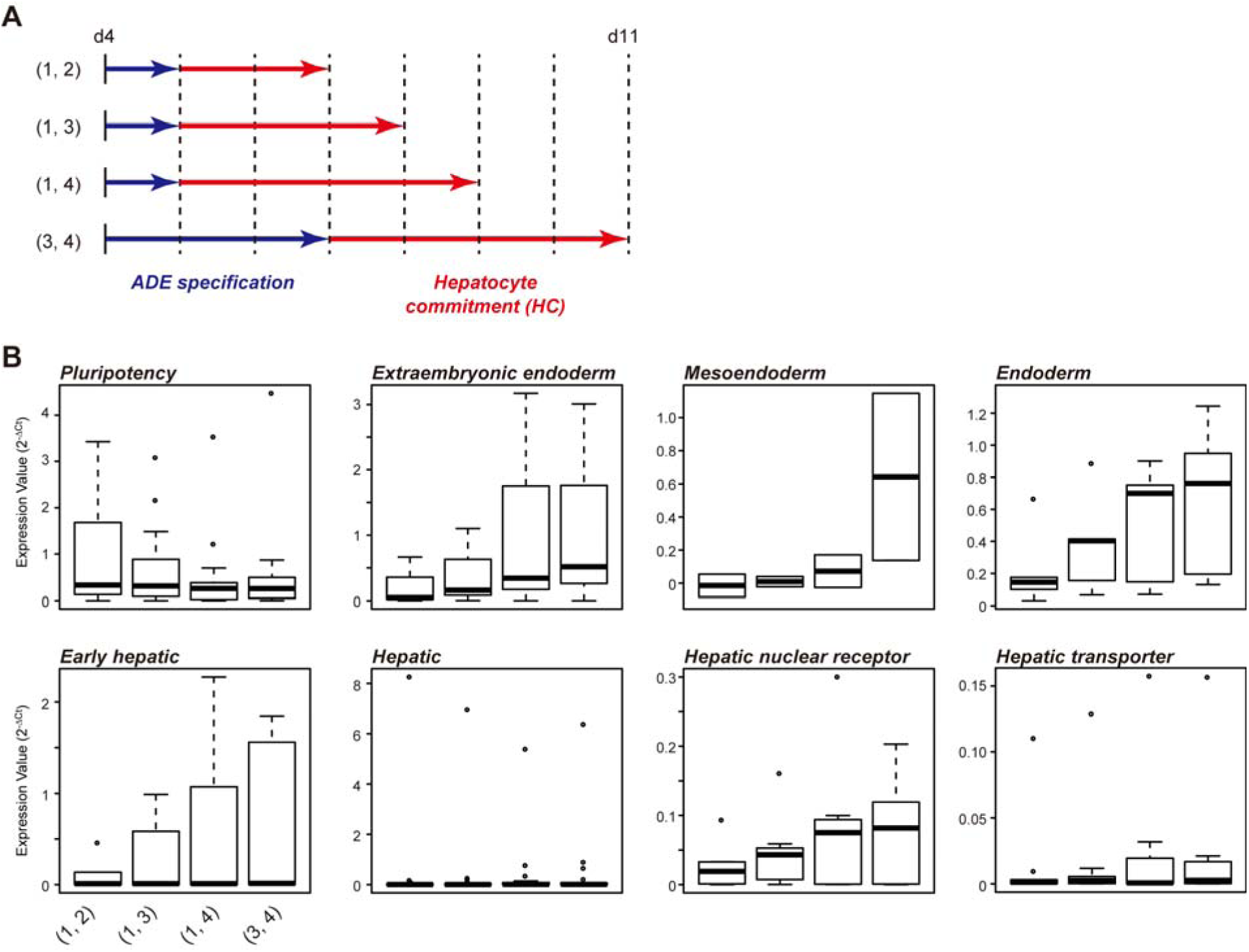
Quantitative RT-PCR analysis reveals optimal periods for anterior definitive endoderm (ADE) specification and hepatocyte commitment. (**A**) Tested periods for ADE specification and hepatocyte commitment. (**B**) Gene-expression patterns of 253G1 hiPSCs treated for ADE specification and hepatocyte commitment. (1, 2) represents 1-day treatment of ADE specification and 2-day treatment for hepatocyte commitment. Center lines show the medians; box limits indicate 25% and 75%; whiskers extend 1.5 times the interquartile range from 25% and 75%. These experiments were carried out at least three times independently.

Following 1 day of treatment for ADE specification, cells continued to express most of the pluripotency associated genes including *NANOG*, *SOX2*, and *OCT4* (or *POU5F1*) (**Fig. 3B**, “Pluripotency”). However, when the treatment for ADE specification was extended to 3 days, the expression levels of pluripotency associated genes was decreased, whereas that of genes associated with extraembryonic endoderm (*SOX7*, *LAMB1*, and *HNF1A*), mesoendoderm (*GSC* and *NODAL*), endoderm (*FOXA2*, *SOX17, CXCR4*, *HNF1B*, and *HNF4A*), and early hepatic cells (*AFP*, *CYP3A7*, *DLK1*, *PROX1*, and *TBX3*) was increased. Although 1-day treatment for ADE specification did not induce expression of genes associated with hepatic nuclear receptors and/or transporters, expression of these genes was observed on days ~2 and ~3. These results indicated that 3 and 4 days of treatment for ADE and HC, respectively, were optimal to obtain hepatic progenitor cells.

### Evaluation of hepatocyte differentiation status by observing gene-expression patterns

To confirm the gene expression associated with hepatic differentiation, we conducted multiplex quantitative RT-PCR, followed by clustering analysis (**Fig. 4, Supplementary Fig. S2** and **Supplementary Table S3**). Clustering analysis based on gene-expression patterns resulted in six clusters. *Class 1* genes in the undifferentiated cells exhibited the highest expression levels, which decreased after the initiation of differentiation. Most *Class1* genes, including *NANOG*, *SOX2*, and *POU5F1*, were associated with pluripotency. *Class 2* included endoderm genes, such as *CXCR4*, *GATA6*, and *SOX17*, which exhibited peak expression at day 5. *Class 3* included early hepatic genes, such as *DLK1* and *TBX3*, which exhibited peak expression at day 12, as well as some genes associated with the cytochrome P450 (CYP) family (*CYP1A2*, *CYP2D6*, and *CYP2E1*). *Class 4* included many hepatic genes, such as *AHSG*, *APOA4*, and *HNF1A*, as well as hepatic transporter genes and solute-carrier-family genes, such as *SLC10A1*, *SLC22A1*, and *SLC22A2*, and exhibited the highest levels of expression at days 20 or 24. *Class 5* genes, including *AQP1*, *B2M*, *CD9, CYP2C8*, *NR3C1*, *SLC22A2*, and *SOX7*, showed the highest expression at day 30, but did not appear to exhibit functional enrichment. *Class 6* showed unique gene-expression patterns, with genes initiating expression on day 20 and peaking at day 35. These includes hepatic genes, such as *ALB*, *SERPINA*, and *TAT*, as well as other early hepatic genes. These gene-expression data suggested that our hepatic differentiation method resulted in a natural developmental process.

**Fig. 4.**
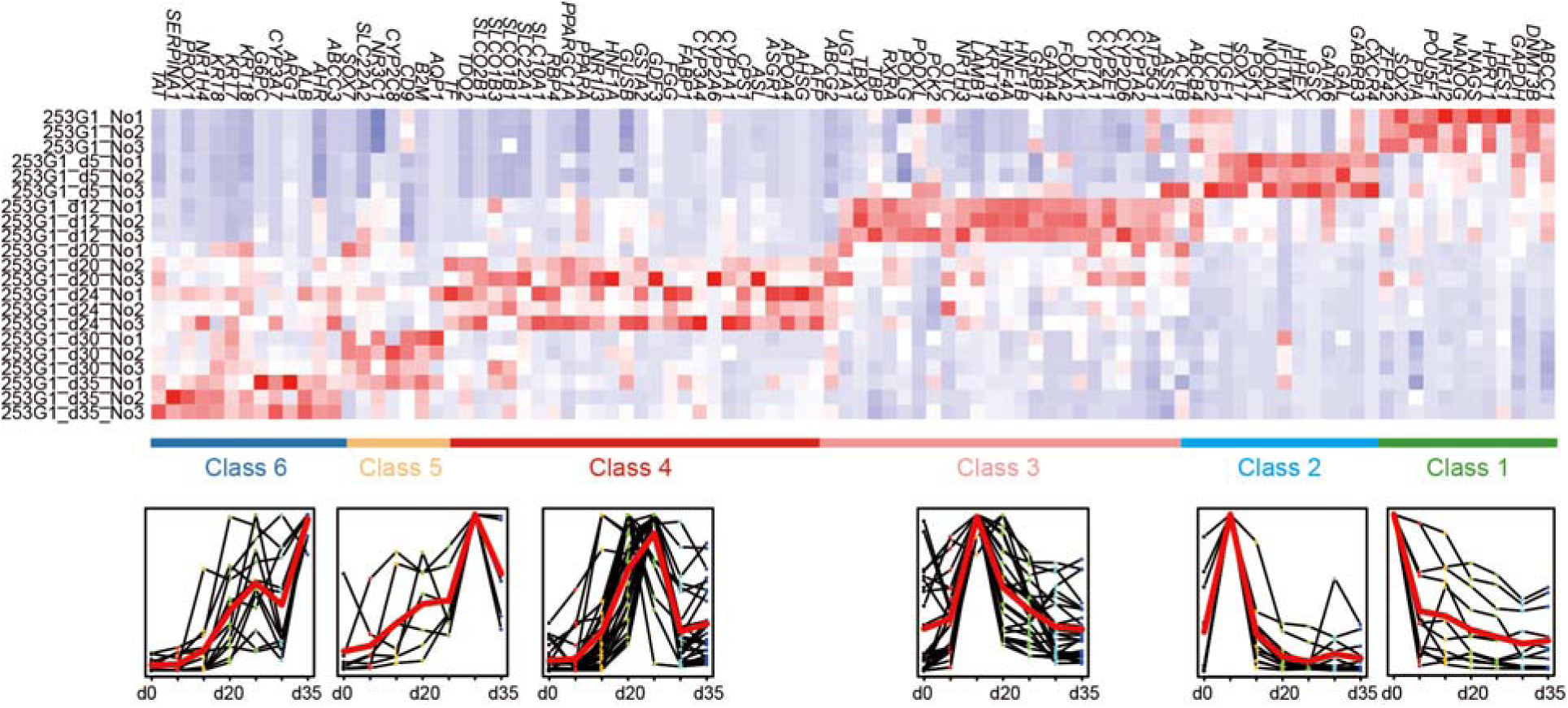
Gene-expression analysis investigating the overall gene-expression signatures from pluripotent status to hepatocyte differentiation. Clustering analysis based on 96 gene expression patterns divided the tested genes into six clusters. An expression trend for each group during the hepatocyte-differentiation process is also plotted.

### Differences in gene expression between hPSC-derived hepatocyte-like cells and liver lysate

To investigate similarities between mature liver and cultured cells, we compared gene-expression patterns between hPSC-derived hepatocyte-like cells and mature liver lysate (**Fig. 5** and **Supplementary Table S3** and **S4**) using hPSC-derived hepatocyte-like cells at days 0, 24, and 35 of differentiation and liver lysate from a healthy donor. Hepatic marker genes, such as *ALB* and *HNF4A*, exhibited significant increases during the culture process, indicating progression of differentiation and increasing similarity between hPSC-derived hepatocyte-like cells and mature liver lysate. Additionally, genes including *AFP*, *APOA4*, *KRT7*, *KRT8*, *KRT18, KRT19, DLK1, GATA6*, and *HNF1B* exhibited increased expression between days 0 and 24, which was higher than the expression of these genes observed in the liver lysate. *AFP* is a marker of hepatoblasts and fetal livers, and *APOA4* exhibits high expression levels in fetal liver. *KRT7*, *KRT8*, *KRT18*, and *KRT19* are expressed in cholangiocytes, *DLK1* is a hepatic progenitor-cell-surface marker, and *GATA6* and *HNF1B* are transcription factors. Specifically, both AFP and DLK1 were strongly expressed in the hPSC-derived cells to a greater degree than in liver lysate, indicating that these cells did not reach the status mature adult liver. However, genes from the CYP family in hPSC-derived cells exhibited lower expression levels relative to those observed in liver lysate, suggesting that our method requires improvement in terms of CYP-family expression.

**Fig. 5.**
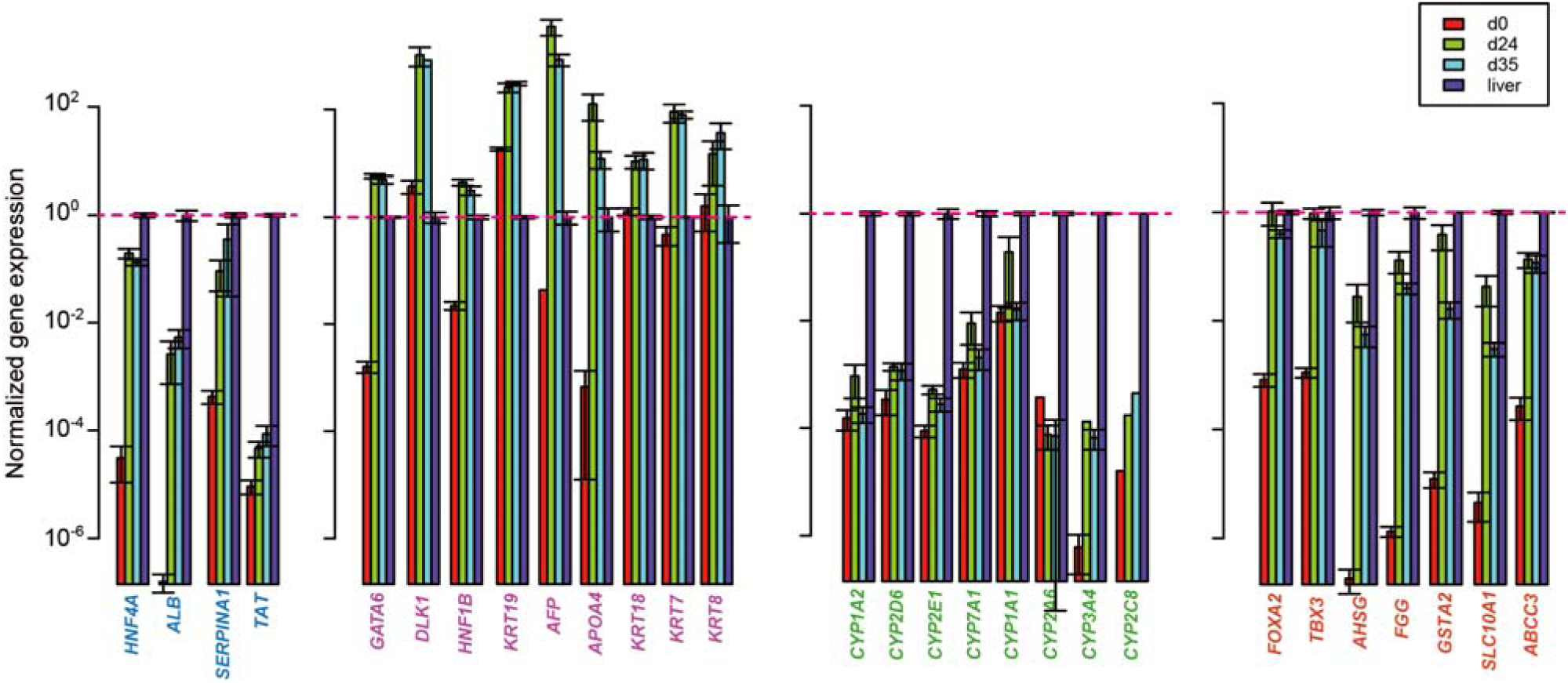
Comparison of gene-expression patterns between hepatocyte-like cells derived from 253G1 hiPSCs and liver lysate. Genes associated with hepatic, hepatoblast, and cytochrome P450 (CYP) family genes were measured by quantitative RT-PCR. Gene-expression values were normalized using the values from liver lysate genes as standards. These experiments were carried out at least three times independently.

### Matured hPSC-derived hepatocyte-like cells in the Liver-on-a-Chip

To facilitate maturation levels for establishment of the Liver-on-a-Chip platform, we developed a simple 3D-culture method using a microfluidic device. We hypothesized that 3D-culture conditions would provide a more suitable environment for hepatocytes as compared with conventional 2D-culture methods^27^ toward mimicking 3D micro liver tissues (**Fig. 6A**). Previously, we reported a microfluidic device enabling 3D culture of hPSCs^29,30^ (**Fig. 6B**), which was utilized in the present study to perform 3D culturing of hepatocyte-like cells from day 12 to enhance the maturation process. The microfluidic device was made of polydimethylsiloxane (PDMS), which exhibits good biocompatibility and gas permeability,^47^ and contained two microfluidic 3D cell-culture chambers [10 mm (L) × 1.5 mm (W) × 150 µm (H)] with a cell inlet (0.75-mm diameter) and a medium reservoir (3-mm diameter). To confirm the maturity of hPSC-derived hepatocyte-like cells at the protein level, we conducted immunocytochemistry for one of the maturation markers, α1 anti-trypsin (A1AT). Our results indicated that the A1AT hepatocyte-maturation marker was more strongly expressed in 3D-cultured hepatocyte-like cells derived from H9 hESCs over 33 days using our method as compared with levels observed in 2D-cultured cells (**Fig. 6C** and **Supplementary Fig. S3**). Although undifferentiated H9 hESCs did not express hepatic genes, we also confirmed increased mRNA expression of other hepatocyte-marker genes, such as *UGT1A1*, *CYP3A4*, *CYP2A9*, *CYC2C19*, and *CYP2D6*, in the 3D-cultured H9 hESC-derived hepatocyte-like cells relative to levels observed in 2D-cultured cells (**Fig. 6D** and **Supplementary Table S5**). In contrast, HepG2 cells did not express *UGT1A1*, *TDO1*, *MDR/TAP*, *CYP3A4*, *CYP2A9*, *CYP2C19*, or *CYP2D6*. As expected, the liver extract from the healthy donor strongly expressed all tested genes at levels exceeding those of hepatocyte-like cells derived from H9 hESCs. These results indicated that the hepatocyte-like cells derived from H9 hESCs exhibited improved gene expression relative to a HepG2 cell line and 2D-cultured hepatocyte-like cells, but were unable to match levels observed in liver extracts from a healthy donor.

**Fig. 6.**
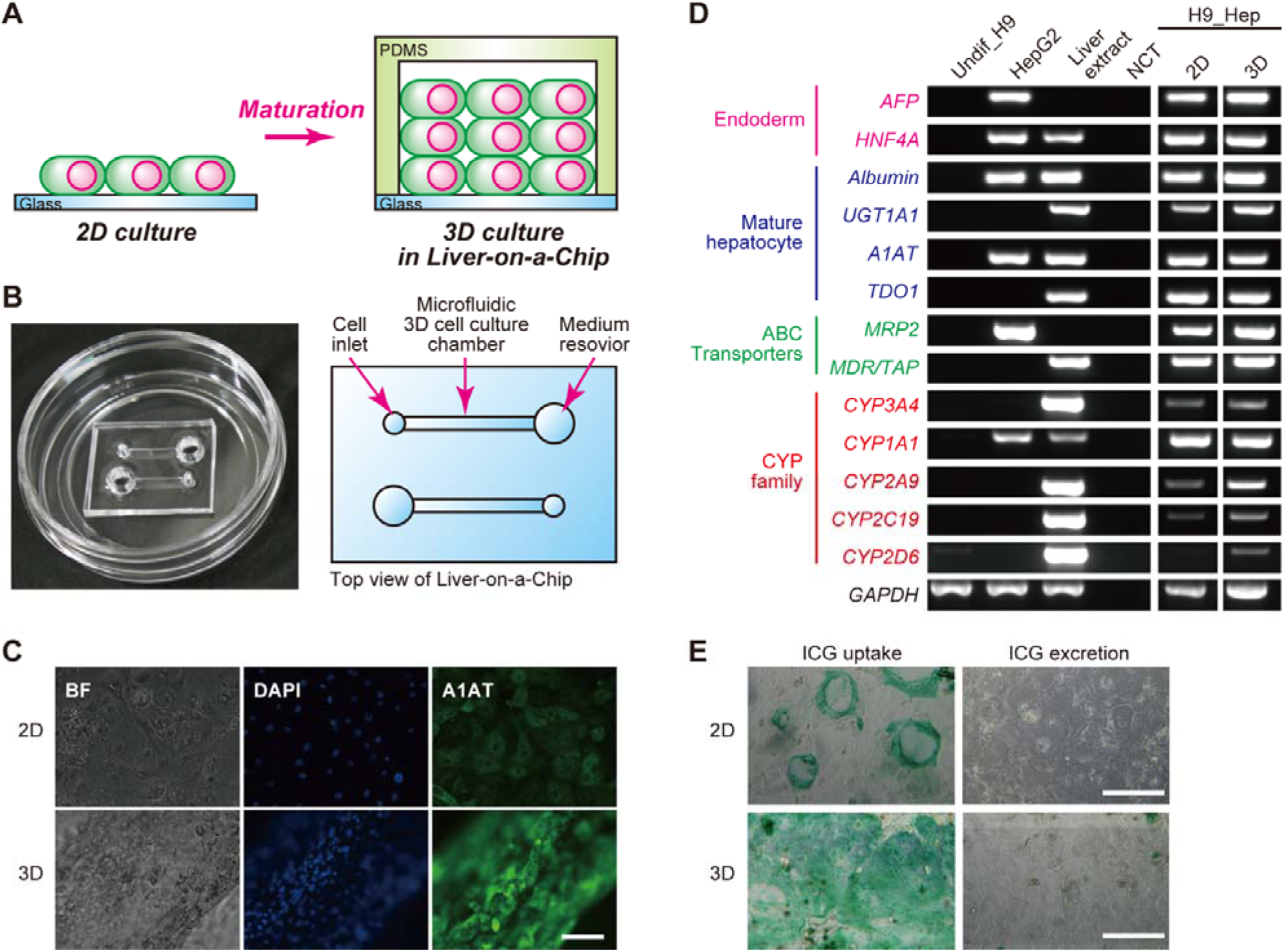
Simple Liver-on-a-Chip platform with matured hPSC-derived hepatocyte-like cells. (**A**) Illustration of 2D and Liver-on-a-Chip 3D cultures for hepatocyte maturation. (**B**) Photograph of a microfluidic 3D-culture device^29,30^ for the Liver-on-a-Chip platform and the structure of a microfluidic cell-culture chamber [10 mm (L) × 1.5 mm (W) × 150 µm (H)] with a cell inlet (0.75-mm diameter) and medium reservoir (3-mm diameter). (**C**) Immunocytochemistry to visualize the hepatocyte-maturation marker α1 anti-trypsin (A1AT) in H9 hESC-derived hepatocyte-like cells. “2D” and “3D” represent hepatocyte-like cells derived from H9 hESCs cultured in a conventional 35-mm culture dish and a microfluidic 3D cell-culture device, respectively. Scale bar represents 20 µ m. (**D**) Representative gel electrophoresis of RT-PCR products for mRNAs associated with the cytochrome P450 family (e.g., *CYP3A4*, *CYP1A1*, *CYP2A9*, *CYP2C19*, and *CYP2D6*), ATP-binding cassette (ABC) transporters (*MRP2* and *MDR/TAP*), hepatocyte-maturation markers (Albumin, *UGT1A1*, *A1AT*, and *TDO1*), and endoderm markers (*AFP* and *HNF4A*), obtained from independently triplicated experiments. HepG2 hepatocellular carcinoma cells were used as a control, and a liver extract from a healthy donor was used as a positive control. H9_Hep and Undif_H9 represent hepatocyte-like cells derived from H9 hESCs and undifferentiated H9 hESCs, respectively. (**E**) Microphotographs of 2D- and microfluidic 3D-hepatocyte-like cells treated with indocyanine green (ICG) for 1 h to visualize ICG uptake and then excretion after 24 h. Scale bar represents 500 µm.

To investigate differences in functionality, an uptake/excretion assay for indocyanine green (ICG), which is a non-invasive marker of drug uptake/excretion in the liver,^45,46^ was performed on cells from both 3D-and 2D cultures (**Fig. 6E** and **Supplementary Fig. S4**). Notably, 3D-cultured cells showed greater ICG uptake within 1 h compared to that of 2D-cultured cells, with most cells also effectively excreting the ICG after 24 h. These results indicated that hPSC-derived hepatocyte-like cells in the Liver-on-a-Chip microfluidic 3D culture device appeared more mature and functional, compared with those of 2D-cultured cells.

## Experimental

**hPSC culture**. hESCs were used according to the guidelines of the ethnical committee of Kyoto University. H9 hESCs were purchased from WiCell Research Institute (Madison, WI, USA). 253G1 hiPSCs were provided by RIKEN BioResource Research Center (Ibaraki, Japan). H9 hESCs and 253G1 hiPSCs were used in this study. Prior to culturing, hESC-certified Matrigel (Corning, Corning, NY, USA) was diluted with Dulbecco’s modified Eagle medium (DMEM)/F12 medium (Sigma-Aldrich, St. Louis, MO, USA) at a 1:75 (v/v) ratio and coated onto a culture dish. Matrigel was incubated in a dish for 24 h at 4°C. Then, excess Matrigel was removed and the coated dish was washed with fresh DMEM/F12 medium.

mTeSR-1-defined medium (Stem Cell Technologies, Vancouver, Canada) was used for daily culturing of hPSCs. For passaging, cells were dissociated with TrypLE Express (Life Technologies, Carlsbad, CA, USA) for 3 min at 37°C and harvested. A cell strainer was used to remove undesired cell aggregates from the cell suspension, and cells were centrifuged at 200 × *g* for 3 min and resuspended in mTeSR-1 medium. Cells were counted using a NucleoCounter NC-200 (Chemetec, Baton Rouge, LA, USA). mTeSR-1 medium containing 10 µM of the ROCK inhibitor Y-27632 (Wako, Osaka, Japan) was used to prevent apoptosis of dissociated hPSCs on day 1. mTeSR-1 medium without the ROCK inhibitor was used on subsequent days, with daily medium changes.

**Hepatic differentiation from hPSCs**. Prior to inducing differentiation, a cell-culture dish was coated with 0.1% gelatin in phosphate-buffered saline (PBS) at 25°C room temperature for 30 min. The gelatin solution was then aspirated and DMEM/F12 medium (Sigma-Aldrich) supplemented with 10% (v/v) fetal bovine serum (FBS), penicillin/streptomycin, and 100 µM α-mercaptoethanol (Sigma-Aldrich) was introduced onto the culture dish for serum coating at 37°C for 24 h. The coated dish was then rinsed with fresh medium.

To induce endoderm differentiation, cultured hPSCs were washed with PBS and treated with TryPLE Express at 37°C for 5 min, followed by the addition of basal medium and transfer of the cell suspension into a 15-mL tube. Cells were centrifuged at 200 × *g* for 3 min and the supernatant was removed. Cells were resuspended in mTeSR-1 medium supplemented with 10 µM Y27632 and 100 ng mL^−1^ activin A, plated on a serum-coated culture dish, and cultured in a humidified incubator at 37°C with 5% CO_2_ for 24 h. At the end of day 1, the medium was replaced with fresh mTeSR-1 medium supplemented with 10 µM Y27632 and 100 ng mL^−1^ activin A and cultured for another 24 h. On day 2, the medium was replaced with mTeSR-1 medium supplemented with 10 µM Y27632, 100 ng mL^−1^ activin A, 10 ng mL^−1^ BMP-4, 10 µM LY294002, and 3 µM CHIR99021, and cells were incubated for 24 h. On day 3, medium was replaced with mTeSR-1 medium supplemented with 10 µM Y27632, 100 ng mL^−1^ activin A, 10 ng mL^−1^ BMP-4, and 10 µM LY294002, and cells were incubated for 24 h. On day 4, medium was replaced with Roswell Park Memorial Institute (RPMI) medium supplemented with B-27 (Life Technologies), 100 ng mL^−1^ activin A, and 100 ng mL^−1^ bFGF, and cells were incubated for 24 h.

To induce ADE specification, cells were treated with RPMI medium supplemented with 50 ng mL^−1^ activin A, with daily medium changes for 3 days. Cells were then treated with RPMI medium supplemented with 20 ng mL^−1^ BMP-4 and 10 ng mL^−1^ FGF-10, with daily medium changes for 4 days.

On day 12, the medium was replaced with hepatocyte basal medium (Lonza, Basel, Switzerland) supplemented with 30 ng mL^−1^ oncostatin M and 50 ng mL^−1^ hepatocyte growth factor to induce maturation of the differentiated hepatocytes. Cells were incubated at 37°C, and medium was changed every 2 days.

**HepG2 cell culture**. HepG2 cells were provided by American Type Culture Collection (Manassas, VA, USA). HepG2 cells were cultured with DMEM supplemented with 10% (v/v) FBS, 1% penicillin/streptomycin, and 1 mM nonessential amino acids, and the medium was changed every 2 to 3 days. The cells were passaged with trypsin-EDTA solutions at a 1:10 to 1:20 subculture ratio.

**Flow cytometry**. Cells were harvested with TrypLE Express and rinsed with PBS twice prior to cell counting. For staining with antibodies, cells were diluted to a final concentration of 1 × 10^7^ cells mL^−1^ in PBS supplemented with 2% fetal calf serum (FCS). Fluorescence-labeled antibodies were added and incubated at room temperature for 30 min. As a negative control, specific isotype controls were used. After removing excess antibodies by centrifugation at 300 × *g* for 5 min, cells were washed with PBS containing 2% FCS, and cell suspensions were applied to a FACS Canto II (BD Biosciences, Franklin Lakes, NJ, USA) for flow cytometric analysis. Data analysis was performed using FlowJo software (v9; FlowJo, LLC, Ashland, OR, USA).

**RT-PCR**. Total RNA was purified using an RNeasy Mini Kit (Qiagen, Hilden, Germany), and 1 µg of total RNA was reverse transcribed to generate cDNA using PrimeScript RT master mix (Perfect Real Time; TaKaRa Bio, Shiga, Japan). A reaction mixture (25 µL) containing 20 ng cDNA, 0.2 µM PCR primers (**Supplementary Table S1**), and 5 U of Taq DNA polymerase (TaKaRa Bio) was subjected to PCR using a thermal cycler (Applied Biosystems 7300 real-time PCR system; Applied Biosystems, Foster City, CA, USA). PCR was performed with 30 to 35 cycles (94°C for 30 s, 58°C for 30 s, and 72°C for 60 s). PCR products (10 μL) were electrophoresed on 1.2% agarose gels and visualized by GelRed nucleic acid staining (Biotium, Fremont, CA, USA).

**Quantitative RT-PCR array**. As a positive control of human normal liver, Human Total Liver RNA was purchased from TaKaRa Bio. Total RNA (2.5 µg) was reverse transcribed to generate cDNA using PrimeScript RT master mix. A reaction mixture (21 µL) containing 20 ng cDNA, 12.5 µL SYBR Premix Ex Taq II (Tli RNaseH Plus; TaKaRa Bio), and 0.5 µL ROX reference dye was introduced into wells of a 96-well plate of Human PrimerArray hepatic differentiation (TaKaRa Bio), according to manufacturer instruction, to assess hepatocyte-specific gene expression (**Supplementary Table S2**). PCR conditions included an initial incubation at 95°C for 30 s, followed by 40 cycles of 95°C for 5 s and then 60°C for 31 s on an Applied Biosystems 7300 real-time PCR system.

**Immunocytochemistry**. Cells were fixed with 4% paraformaldehyde in PBS for 20 min at 25°C and then permeabilized with 0.5% Triton X-100 in PBS for 16 h at 25°C. Subsequently, cells were blocked in PBS (5% normal goat serum, 5% normal donkey serum, 3% bovine serum albumin, 0.1% Tween-20) at 4°C for 16 h and then incubated at 4°C for 16 h with the primary antibody [anti-human A1AT rabbit IgG, 1:800; Dako, Glostrup, Denmark] in PBS with 0.5% Triton X-100. Cells were then incubated at 37°C for 60 min with a secondary antibody (AlexaFluor 488 Donkey anti-rabbit IgG, 1:1000; Jackson ImmunoResearch, West Grove, PA, USA) in blocking buffer prior to a final incubation with 4′,6-diamidino-2-phenylindole (DAPI) at 25°C for 30 min.

**Image acquisition**. The sample containing cells was placed on the stage of a Nikon ECLIPSE Ti inverted fluorescence microscope equipped with a CFI plan fluor 10×/0.30 N.A. objective lens (Nikon, Tokyo, Japan), CCD camera (ORCA-R2; Hamamatsu Photonics, Hamamatsu City, Japan), mercury lamp (Intensilight; Nikon), XYZ automated stage (Ti-S-ER motorized stage with encoders; Nikon), and filter cubes for fluorescence channels (DAPI and GFP HYQ; Nikon). For image acquisition, the exposure times were set at 500 ms for DAPI and 500 ms for GFP HYQ (for A1AT).

**ICG uptake/excretion assay**. Briefly, 1 mg mL^−1^ ICG was dissolved in hepatocyte-maturation medium and cells were treated with ICG solution for 1 h, rinsed with hepatocyte-maturation medium, and then observed using a bright-field microscope (Olympus, Tokyo, Japan). After 24 h, cells were observed again to visualize excretion capability.

**Microfluidic device fabrication**. A microfluidic device was fabricated using stereolithographic 3D-printing techniques and solution cast-molding processes.^29^ The mold for the microfluidic channels was produced using a 3D printer (Keyence Corporation, Osaka, Japan). Sylgard 184 PDMS two-part elastomer (10:1 ratio of pre-polymer to curing agent; Dow Corning Corporation, Midland, MI, USA) was mixed, poured into a 3D-printed mold to produce a 5-mm-thick PDMS layer, and de-gassed by using a vacuum desiccator. The PDMS material was then cured in an oven at 65°C for 48 h. After curing, the PDMS form was removed from the mold, trimmed, and cleaned. The PDMS form and a glass dish or plastic plate were corona-plasma-treated (Kasuga Denki, Inc., Kawasaki, Japan) and bonded together by baking in an oven at 80°C.

**Preparation of the Liver-on-a-Chip microfluidic 3D culture device**. Prior to use, a microfluidic 3D cell-culture device was sterilized in 70% ethanol and placed under ultraviolet light in a biosafety cabinet for 30 min. A microfluidic cell-culture chamber was then coated with Corning Matrigel hESC-qualified matrix (Corning) diluted to 1:75 (v/v) with DMEM/F12 (Sigma-Aldrich). After a 1-day incubation at 4°C, excess Matrigel was removed, and the coated dish was washed with fresh DMEM/F12. Cells were harvested using trypsin and collected in a 15-mL tube. Following centrifugation, cells were suspended in hepatocyte-maturation medium and introduced into a microfluidic device via a cell inlet. The microfluidic device was placed in a humidified incubator at 37°C with 5% CO_2_ atmosphere, and the medium was changed daily.

**Statistical analysis of gene expression**. A two-tailed Student’s *t*-test was carried out in Microsoft Excel. For gene expression analysis, gene clusters were generated using normalized gene-expression values for the quantitative RT-PCR array by the Consensus Cluster algorithm in the *R* package “ConsensusClusterPlus”.^48^ Mean values were calculated in each cluster and for each day, and fold changes were calculated using liver values as a standard.

## Conclusions

In this study, we developed a Liver-on-a-Chip 3D culture platform with matured hPSC-derived hepatocyte-like cells. In particular, we firstly established an efficient and robust method to induce hPSC differentiation into functional hepatocyte-like cells. By optimizing the basal medium, combination of chemicals, and initial cell-seeding density for endoderm differentiation, we obtained a homogeneous population of endoderm-differentiated cells highly expressing CXCR4 from dissociated hPSCs. Additionally, we optimized the periods required for ADE specification and HC according to quantitative PCR analysis. To our knowledge, this represents the first demonstration of application of a microfluidic 3D cell-culture platform for the maturation of hepatocyte-like cells.

Liver-on-a-Chip platforms offer critical opportunities for drug screening and chemical-safety testing in a variety of industries including cosmetics and agriculture. However, there are limited numbers of cell sources (e.g., primary hepatocytes) available, with the number and quality varying by donor. Although current hepatocyte cell lines, such as HepG2, have been used as alternatives, they often show different characteristics from primary hepatocytes. In addition, current 2D-culture conditions in a chip do not allow the expression of proper hepatic functions. In this study, we successfully developed a Liver-on-a-Chip platform by establishing a method to induce hPSC differentiation into mature hepatocytes with microfluidic 3D culture to prepare functional hepatocyte-like cells on a chip.

Liver-on-a-Chip platforms may be used for either “High-content analysis (HCA)” or “High-throughput screening (HTS)”. Although a number of Liver-on-a-Chip platforms have been previously reported, these often provided less information than that of current HCA, are exceedingly complicated for general users, and not applicable for HTS. For drug screening and chemical-safety testing platforms, it is necessary to decide which application would be most suitable at the beginning of device development. Our device and cell culture chamber are very simple and offer easy handling without requiring any special instruments. They can be applicable for increased throughput by means of straightforward device format design, such as the use of microplates, following the recommendations of the recommendation of the Society for Biomolecular Screening (SBS). Accordingly, we consider that our Liver-on-a-Chip platform will fulfill current requirements and serve as a useful tool in the field of drug discovery.

## Acknowledgements

Funding was generously provided by the Japan Society for the Promotion of Science (JSPS; 24656502, 26560209, 16K14660, and 17H02083). Funding was also provided by the Terumo Life Science Foundation and Japan Agency for Medical Research and Development. The WPI-iCeMS is supported by the World Premier International Research Centre Initiative (WPI), MEXT, Japan.

